# The diet and feeding behaviour of the black-and-white colobus (*Colobus guereza*) in the Kalinzu Forest, Uganda

**DOI:** 10.1101/832451

**Authors:** Ikki Matsuda, Hiroshi Ihobe, Yasuko Tashiro, Takakazu Yumoto, Deborah Baranga, Chie Hashimoto

## Abstract

One of the goals for primate feeding ecology is to understand the factors that affect inter- and intra-specific variations. Therefore, a detailed description of basic feeding ecology in as many populations and sites as possible is necessary and warrants further understanding. The black-and-white colobus (*Colobus guereza*) or guereza is widely distributed in Africa and is one of the well-studied colobines in terms of their feeding; they demonstrate considerable variation in their diets in response to local conditions. We studied the diet of a group of guerezas in the Kalinzu Forest, Uganda, for over 30 consecutive months using behavioural observation (4,308 h total), phenology, and vegetation surveys. A total of 31 plant species were consumed by the study group. This study group was predominantly folivorous; the majority of their feeding time was involved in feeding on young leaves (87%). However, during certain times of the year, fruits and seeds accounted for 45% of monthly feeding time. Young leaves of *Celtis durandii* were by far the most important food, which constituted 58% of the total feeding records. There was a significant increase in the consumption of fruits and flowers once young leaf availability was low, but their consumption of fruits did not significantly increase even when fruit availability was high. Their monthly dietary diversity increased as the number of available plants with young leaves declined, suggesting that much of the dietary diversity in the study group may be attributable to the young leaf portion of their diet. Our findings may help contribute to a better understanding of the dietary adaptations and feeding ecology of guerezas in response to local environmental conditions.

## Introduction

A considerable amount of knowledge has been accumulated over the past few decades with respect to the dietary information of non-human primates. This has not only served to increase researchers’ knowledge regarding the diet of primates but also contributed in understanding their dietary flexibility in response to the environment (Campbell et al. 2011). Such flexible variations in diet serve to satisfy their nutritional requirements in habitats with seasonally fluctuating food availability, which is influenced by various environmental factors, such as temperature, rainfall, and plant phenology (Hanya et al. 2013; Hemingway and Bynum 2005). Thus, an assessment of dietary variation in relation to seasonal conditions is important for evaluating primate foraging strategies.

Colobine monkeys, which include at least 30 species grouped into 4–9 genera in Asia and Africa (Oates et al. 1994), feed on difficult-to-process foods, including leaves, seeds, and unripe fruits, which they process in their complex, multi-chambered stomachs, wherein bacteria detoxify chemicals from defensive plants and digest cellulose (Chivers 1994). Previously it was assumed that colobine monkeys primarily exploit ubiquitous food sources like leaves because of their foregut-fermentation digestive system; however, there is evidence of considerable variation in their diet (in some cases, they rely heavily on fruits) (Fashing 2011; Kirkpatrick 2011). Particularly, colobines in clades with tripartite stomachs (without praesaccus) tend to rely more on fruits and seeds for food during times when fruits are readily availability (Matsuda et al. 2019).

Foregut-fermenting black-and-white colobuses, aka guerezas (*Colobus guereza*), with tripartite stomachs, which is the species of the genus most widely distributed in Africa (Zinner et al. 2019), have been reported to have considerably varied diets. Some populations have been documented as highly folivorous (Clutton-Brock 1975; Harris and Chapman 2007; Hussein et al. 2017; Oates 1977; Wasserman and Chapman 2003), whereas others eat large quantities of fruit (Fashing 2001; Poulsen et al. 2002). Because one of the goals of feeding ecology is to understand the factors that affect such intra-specific dietary variations across different populations and sites, a detailed description of as many populations and sites as possible is imperative to further this understanding.

We studied the diet of guerezas in the Kalinzu Forest, Uganda, for over 30 consecutive months. We aim to describe the dietary changes among its different species in response to seasonal fluctuations and availability of certain food, including leaves, flowers and fruits. This study presents the systematic information regarding the feeding habits and monthly dietary variation of guerezas in this study site.

## Methods

### Study site and focal group

Observations were conducted from November 2013 to April 2016 in a moist, medium-altitude evergreen forest in the Kalinzu Forest in western Uganda, covering an area of 137 km^2^ (30°07’ E, 0°17’ S; altitude 1,000–1,500 m above sea level) (Hashimoto et al. 1999). From November 2013 to April 2016, the mean minimum and maximum daily temperatures were approximately 14.0°C (SD: 1.9) and 27.2°C (SD: 2.1), respectively. The annual precipitation at the site in 2015 was 1370 mm.

From November 2011 to October 2012, preliminary observations were performed on several guereza groups before the most habituated group was chosen for the study. During these preliminary observations, members of a focal group were identified by describing their individual physical characteristics. At the end of the preliminary observations, the study group included 11 individuals: one alpha male, three adult females, two subadult females, two juveniles, and three infants.

### Behavioural data collection

We observed the habituated focal group for 10–22 d/mo from approximately 7:30 to 16:00. During the observation periods, we conducted scan sampling at 10-min intervals. We recorded the activity (feeding, moving, and resting) of all visible adults and subadults ranging from 1–7 individuals with a mean number of 4.8 ± SD 1.2 individuals per scan. We recorded the food category and collected samples for later identification when they were feeding. We collected data for 30 months from November 2013 to April 2016, except when data collection was interrupted by a field accident in November 2014. The total observation time was 4,308 h, and the monthly observation time was 69–185 h (mean: 139 ± SD 34 h). The observation time per day was 8.00 ± SD 1.01 h.

### Vegetational and phenological survey

In addition to the ten parallel transects, each of which was 5 km in length and built no earlier than 1997 (Hashimoto et al. 1999), to assist in observation and focal following, a trails system was set up in the study site. Based on the ranging data of the focal group during the preliminary observation time, we selected 12 trails that were 180–900 m long (total 4,700 m). We labeled trees ≥10 cm in diameter at breast height (DBH) and vines ≥5 cm in diameter located ≤1.5 m from the trail; hence, the labeled width was 3 m, and the vegetation survey area covered 1.41 ha (3 m × 4,700 m). All labeled trees and vines were taxonomically identified from the Makerere University Herbarium. At the end of the monthly survey, the phenology of the 996 labeled plants along the 12 trails was recorded by examining each plant for the presence or absence of fruits (both ripe and unripe) and flowers (including floral buds). We also recorded the presence or absence of young leaves but used their abundance score, i.e., 1: < 20% of young leaves in a tree, 2: 20% < 50%, and 3: 50% ≤ 100%.

### Data analysis

Data were tested for normality using the Kolmogorov–Smirnov test. To test the correlations between the monthly availability of each plant part (young leaves, fruits, and flowers) and feeding activity on each plant part, data were measured using either the parametric Pearson correlation coefficient for variables that were normally distributed or the nonparametric Spearman’s rank correlation coefficient for variables that were not normally distributed. We used the Holm sequential Bonferroni procedure to correct for multiple comparisons (Holm 1979) and applied the corrected *P*-values. The Shannon–Wiener index of diversity (*H*’) (Pielou 1966) was used to calculate the dietary diversity of each month. The linear mixed model was used to examine whether the monthly dietary diversity of guerezas (response variable) was affected by the availability of leaves, fruits, and/or flowers (explanatory variables). The variance inflation factors were smaller than the cut-off value, i.e., 10 (Quinn and Keough 2002). The Akaike information criterion for small samples (AICc value: Burnham and Anderson 2002) was used to find a good-fitting model (ΔAICc < 2, i.e., the AICc in a model is less than two units larger than in the best model). We used R 3.53 for all the statistical analyses (R-Core-Development-Team 2019), and the significance level was set at 0.05.

## Results

### Vegetation and phenology

We marked 969 trees and 27 vines (68 species, 57 genera, 35 families) along our 12 trails (Table 1). The five most abundant families were Apocynaceae (20.7% of all plants), Meliaceae (12.7%), Oleaceae (10.1%), Rubiaceae (9.4%), and Cannabaceae (9.0%). The five most abundant plant species were *Funtumia africana* (18.5% of all trees), *Carapa procera* (12.0%), *Strombosia scheffleri* (10.1%), *Celtis durandii* (8.0%), and *Musanga leo-errerae* (5.4%). The vegetation survey area covered 1.41 ha (3 m × 4,700 m). The cumulative number of plant species did not appear to reach an asymptote (Fig.1), probably because of the high species diversity in the tropical secondary forest. However, the total number of plant species used as a food source by the study group (39 plant species) was lower than that described in the vegetation survey, indicating that the results of vegetation survey represent the composition of the available plant species by the study group.

**Table 1.**
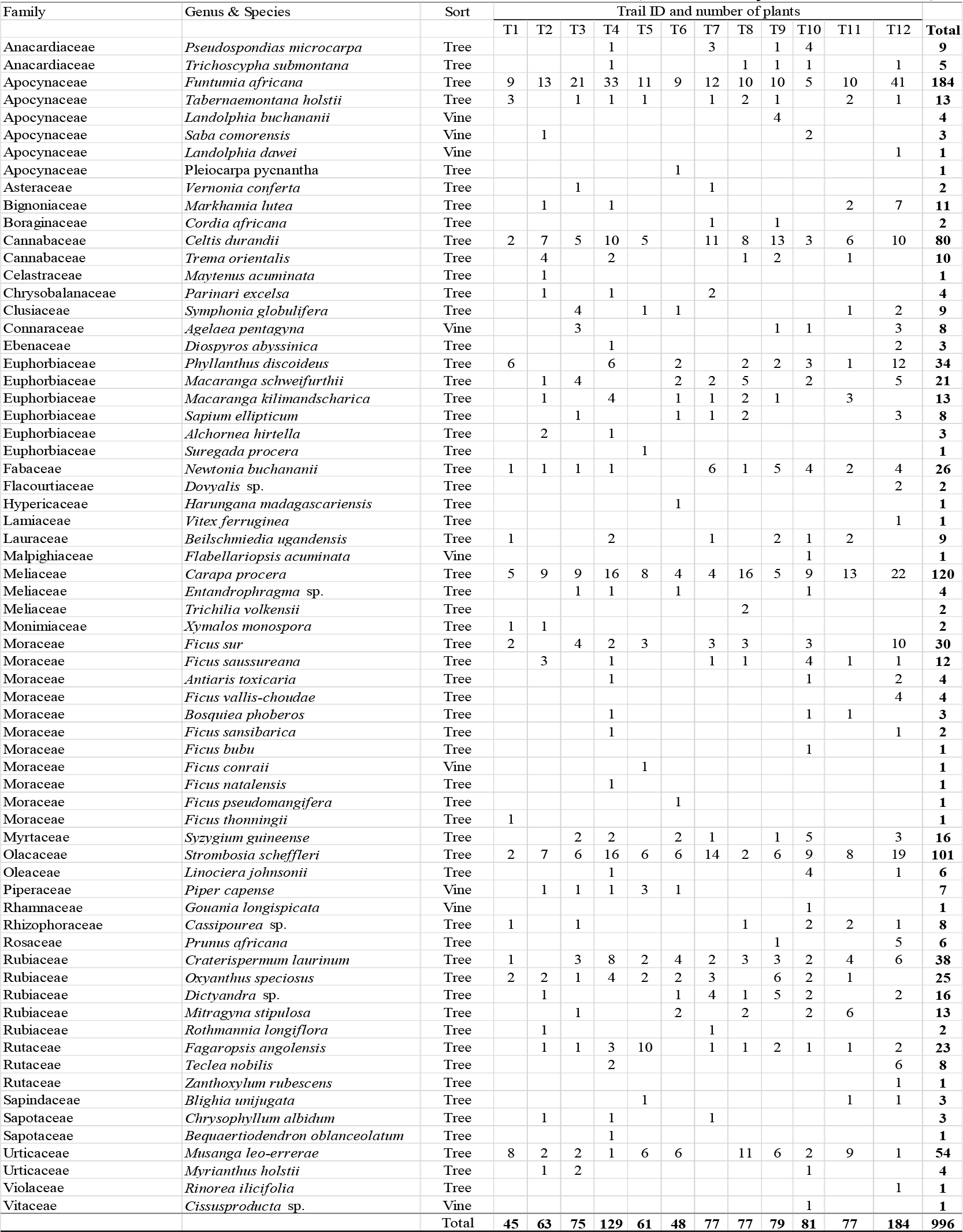
Floral composition along the 12 trails. The total distance of the trails was 4.7 km, and the vegetation survey was conducted for trees with a DBH of ≥10 cm and vines with a diameter of ≥5 cm, located ≤1.5 m from the trail (total survey area = 1.41 ha).

**Figure 1.**
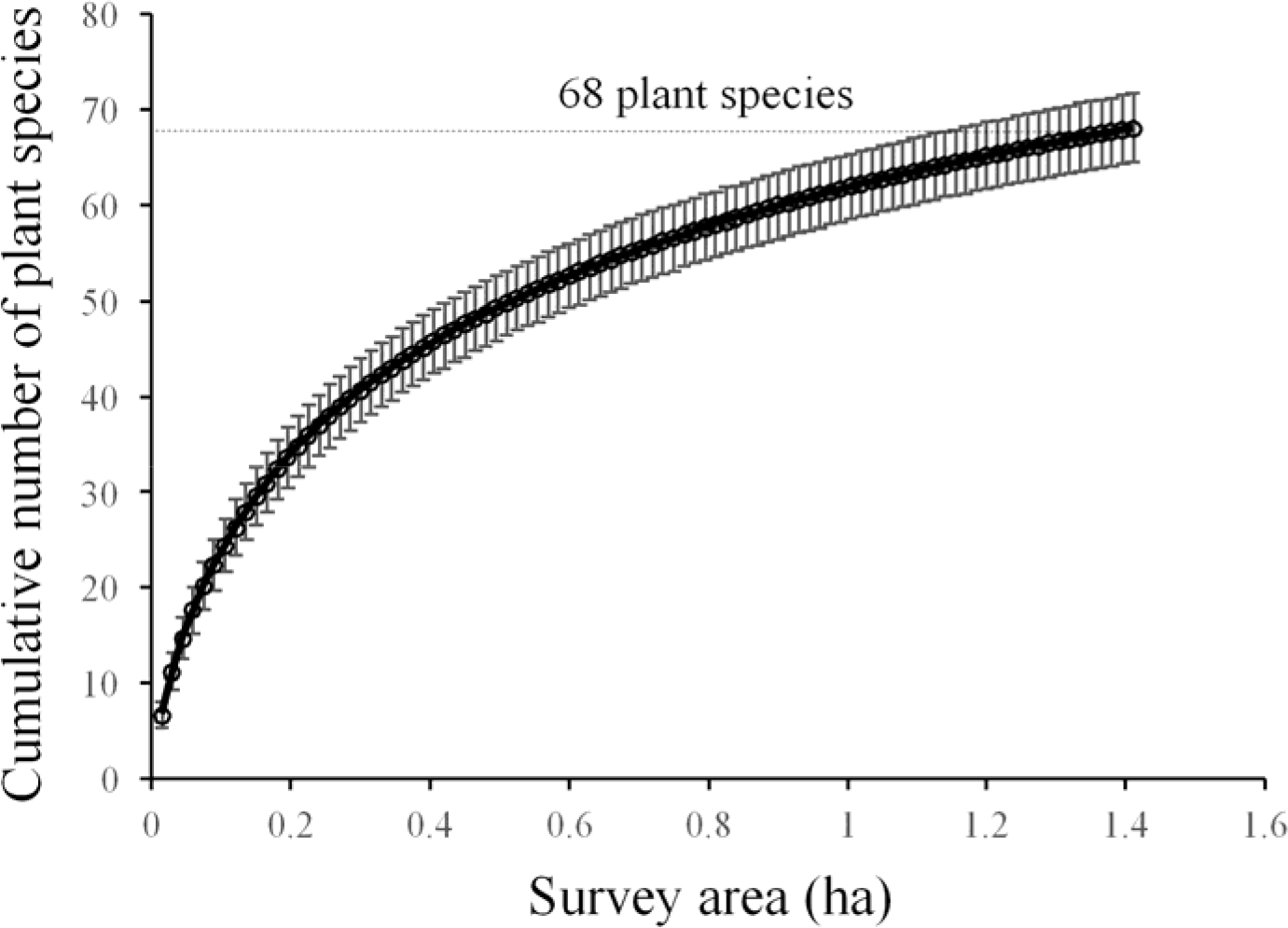
Species accumulation curves (species vs. area curves) of the study site covering the survey area of 1.41 ha. Error bars indicate standard deviation. We performed 100 randomizations using the data of the number of tree species with a DBH of ≥10 cm and vine species with a diameter of ≥5 cm at subplots of 3 m × 50 m and used EstimateS ver. 9 (Colwell 2013) to produce the curves.

Of the 969 trees and vines along the trails, 515 (53.3%) were plant species that we observed the study group to use as a food sources. This included 505 plants with young leaves (52.2%), 392 with fruits (41.0%), and 148 with flowers (15.3%). Food availability, determined as the number of young leafing (but applied scored numbers: see in Methods), fruiting, and flowering potential food plants in the monthly phenological survey, fluctuated seasonally with several peaks within a year (Fig. 2). Young leaves were generally available during all seasons and were more abundant than fruits and flowers.

**Figure 2.**
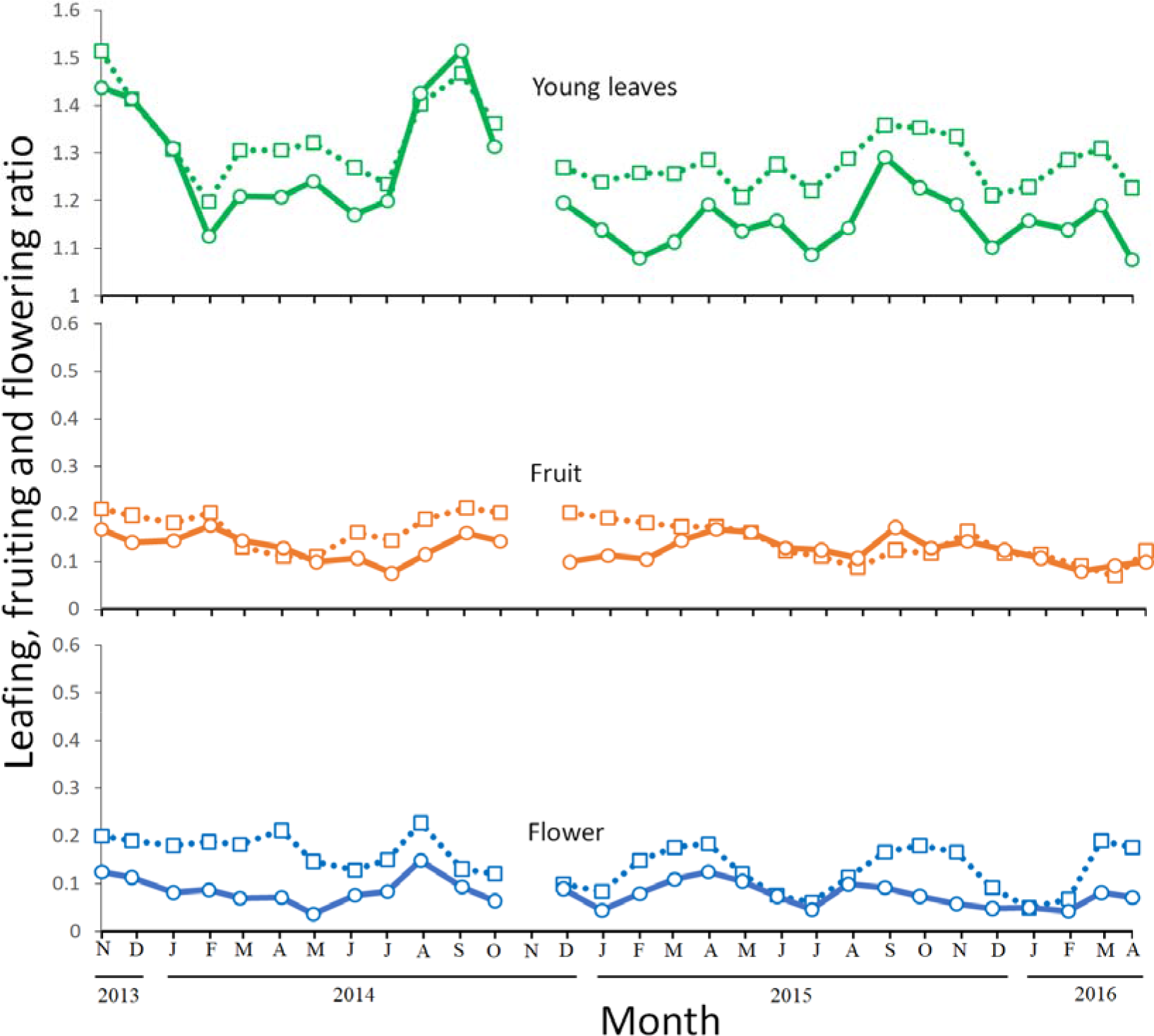
Young leafing, fruiting, and flowering phenology between November 2013 and April 2016. Values indicate ratios of the monthly number of counted plants to young leaves, fruits and flowers; the presence or absence of observed fruits and flowers was examined for each plant, but the presence or absence of young leaves was examined using abundance scores (see Methods section). Square = all plants; circle = guereza’s food species only.

### Overall food habits

The study group frequently consumed young leaves, fruits and flowers; they occasionally consumed mature leaves and other food types including the tree bark of *Carapa procera* and *Eucalyptus grandis* and soil on the ground. The total number of plant species consumed by the group of guerezas during 4,308 h of observation was 31, including two unidentified species (26 genera, 24 families; Table 2); it was 39 plant species including the preliminary observation time. A number of plant species that provided leaves, fruits/seeds, and flowers were 31, 12, and 6, respectively. Overall, the study group fed on young leaves (87.0% of feeding record); fruits (9.8%), including seeds (5.0%), both seeds and pulp of ripe (4.5%) and unripe fruits (0.3%); flowers (1.1%); bark (0.9%); soil (0.8%); and other foods (unspecified foods and mature leaf: 0.4%). Young leaves of *Celtis durandii* (Cannabaceae) were by far the most important food, which constituted 58.1% of the total feeding records, followed by the young leaves of *Prunus africana* (Rosaceae: 11.0%) and the fruits including seeds of *C. durandii* (4.8%).

**Table 2.**
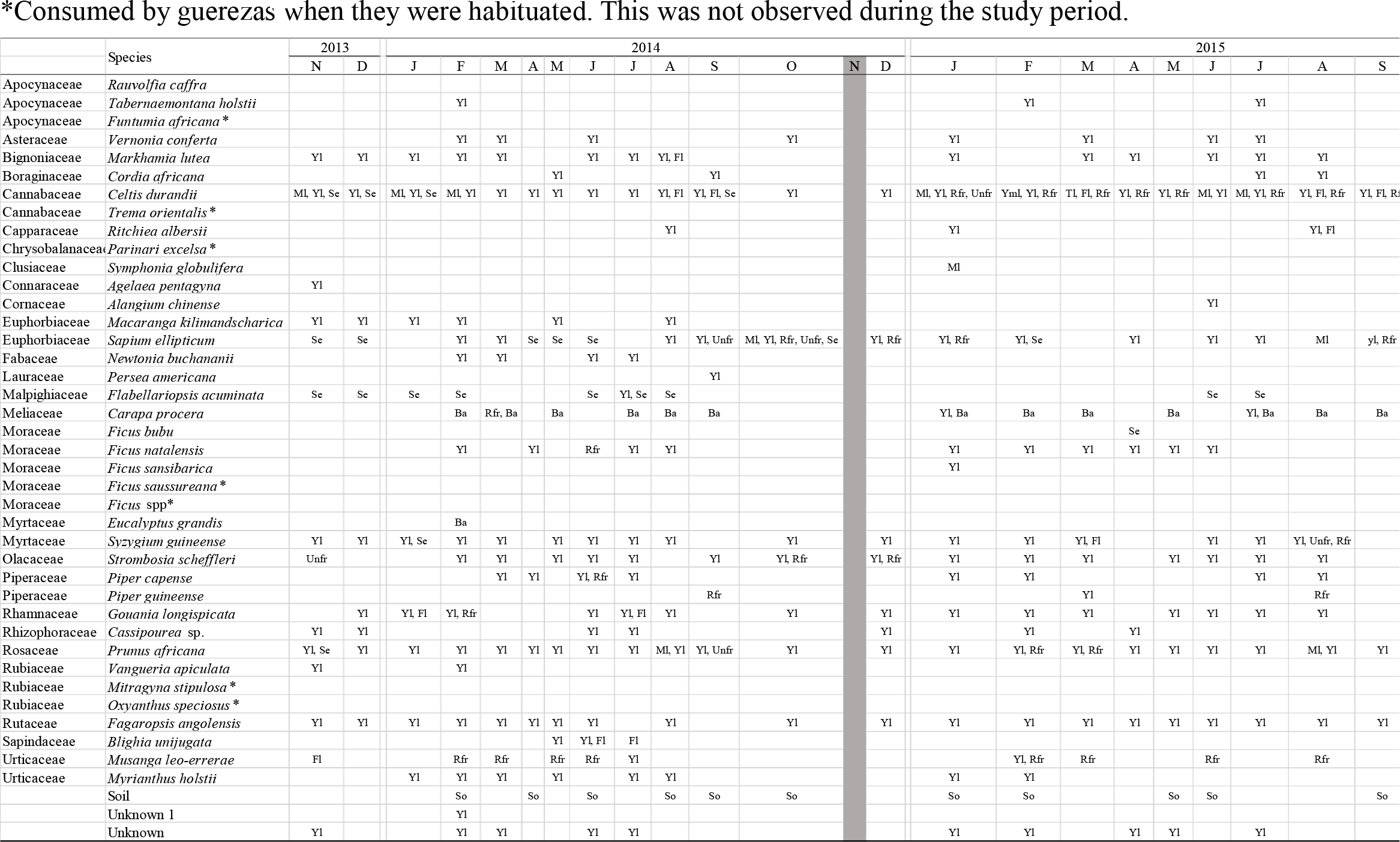
Food items and parts of each item consumed by the study group from November 2013 to April 2016.

### Monthly variation in diet and dietary diversity

We analyzed the seasonal changes in diet to examine the effects of temporal changes in the availability of fruits, flowers and young leaves on the feeding patterns of guerezas, seasonal changes in diet were analyzed (Fig. 3). We observed that the majority of the feeding time in the monkeys of the study group was dedicated to the feeding of young leaves throughout the study period; however, during certain times of the year, fruits/seeds accounted for over 45% of monthly feeding record. Monthly fruit availability was not significantly correlated with fruit-eating (r = 0.42, *P* = 0.069), flower-eating (r = 0.42, *P* = 0.052) and leaf-eating (r = 0.32, *P* = 0.086) activities. Alternatively, monthly flower availability was significantly correlated with flower-eating activity (rs = 0.57, *P* = 0.004) but not with fruit-eating (rs = 0.084, *P* = 0.665) and leaf-eating (rs = 0.31, *P* = 0.213) activities. In addition, monthly leaf availability was significantly correlated with fruit-eating (r = −0.42, *P* = 0.044) and flower-eating (r = −0.52, *P* = 0.011) activities but not with leaf activity (r = −0.32, *P* = 0.089).

**Figure 3.**
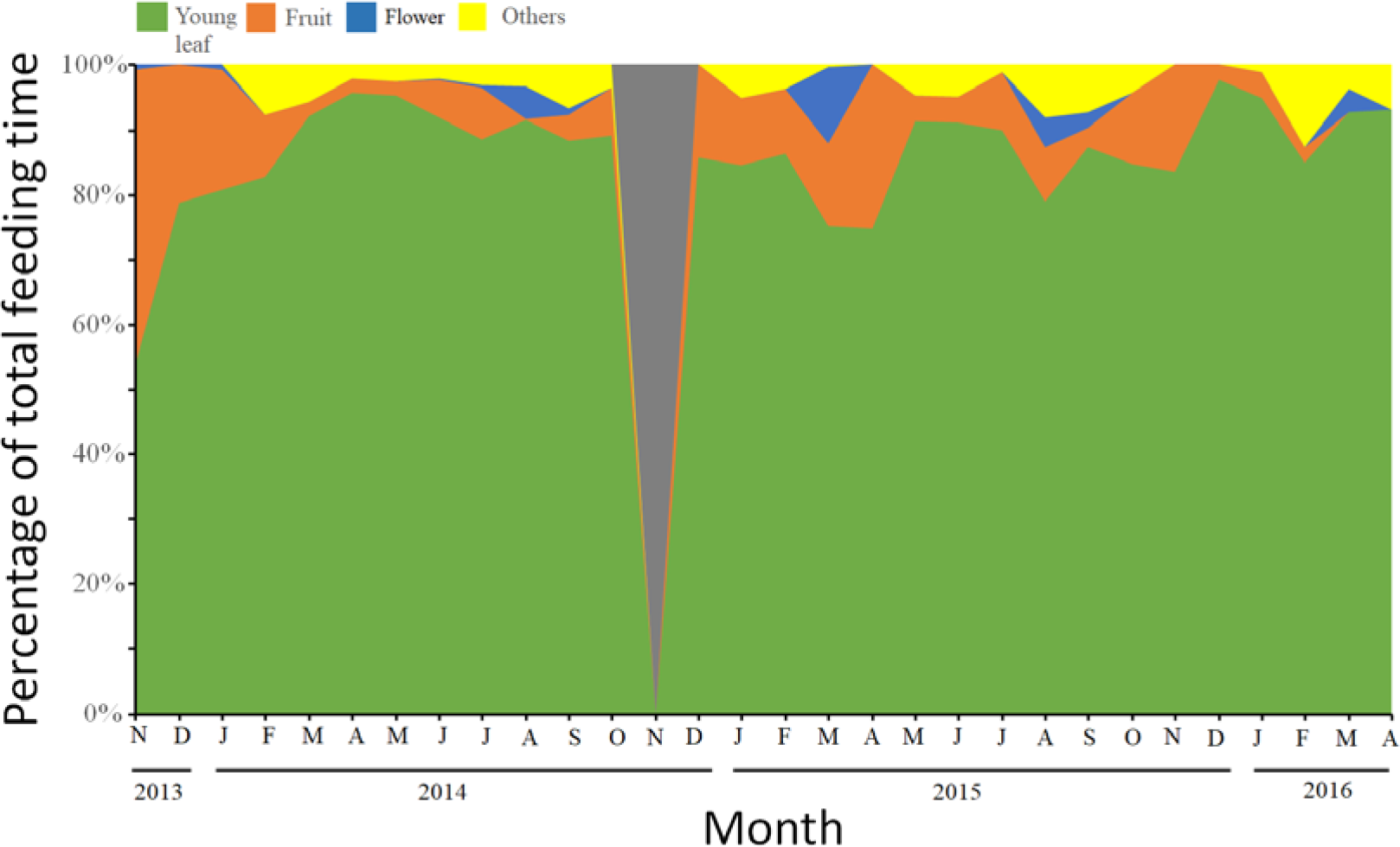
Seasonal changes in the diet composition of the study group; values indicate the percentage of monthly feeding time spent on each food category.

We observed that the young leaves of *C. durandii* were the most important food source for the study group in our site, and its feeding record on young leaves was much higher than that of the second and third most important food sources. The study group consumed *C. durandii* young leaves in each of the study months, ranging from 34.2% in February 2015 to 90.8% in April 2014. The monthly feeding record of *C. durandii* was not significantly correlated with monthly fruit (r = −0.15, *P* = 0.866), flower (r = −0.08, *P* = 0.692) and leaf availabilities (r = −0.28, P = 0.840).

The mean of the 29 monthly Shannon–Wiener indices of food species diversity (*H’*) was 1.37 (range: from 0.35 in April 2015 to 2.19 in February 2014). The best-fit model to explain the monthly dietary diversity included the leaf availability, and the second-best model was the null model (Table 3); the dietary diversity increased with the decreasing leaf availability (Fig. 4).

**Table 3.** Summary of model selection using linear models to investigate whether the dietary diversity (*H’*) of the study group was affected by the availability of fruits, flowers, and young leaves.

| Intercept | Fruit<br>availability | Flower<br>availability | Young leaf<br>availability | df | Log-<br>likelihood | AICc | Δ-AICc | Weight |
| --- | --- | --- | --- | --- | --- | --- | --- | --- |
| 2.7 |  |  | -1.1 | 3.0 | -13.8 | 34.5 | 0.0 | 0.3 |
| 1.4 |  |  |  | 2.0 | -15.2 | 34.9 | 0.4 | 0.3 |
| 2.9 |  | 2.9 | -1.4 | 4.0 | -13.3 | 36.3 | 1.8 | 0.1 |
| 2.7 | 2.4 |  | -1.3 | 4.0 | -13.4 | 36.5 | 2.0 | 0.1 |
| 1.3 | 0.4 |  |  | 3.0 | -15.2 | 37.3 | 2.8 | 0.1 |
| 1.4 |  | -0.1 |  | 3.0 | -15.2 | 37.3 | 2.9 | 0.1 |
| 2.8 | 1.7 | 2.3 | -1.5 | 5.0 | -13.1 | 38.9 | 4.4 | 0.0 |
| 1.3 | 0.6 | -0.3 |  | 4.0 | -15.2 | 40.0 | 5.5 | 0.0 |

**Figure 4.**
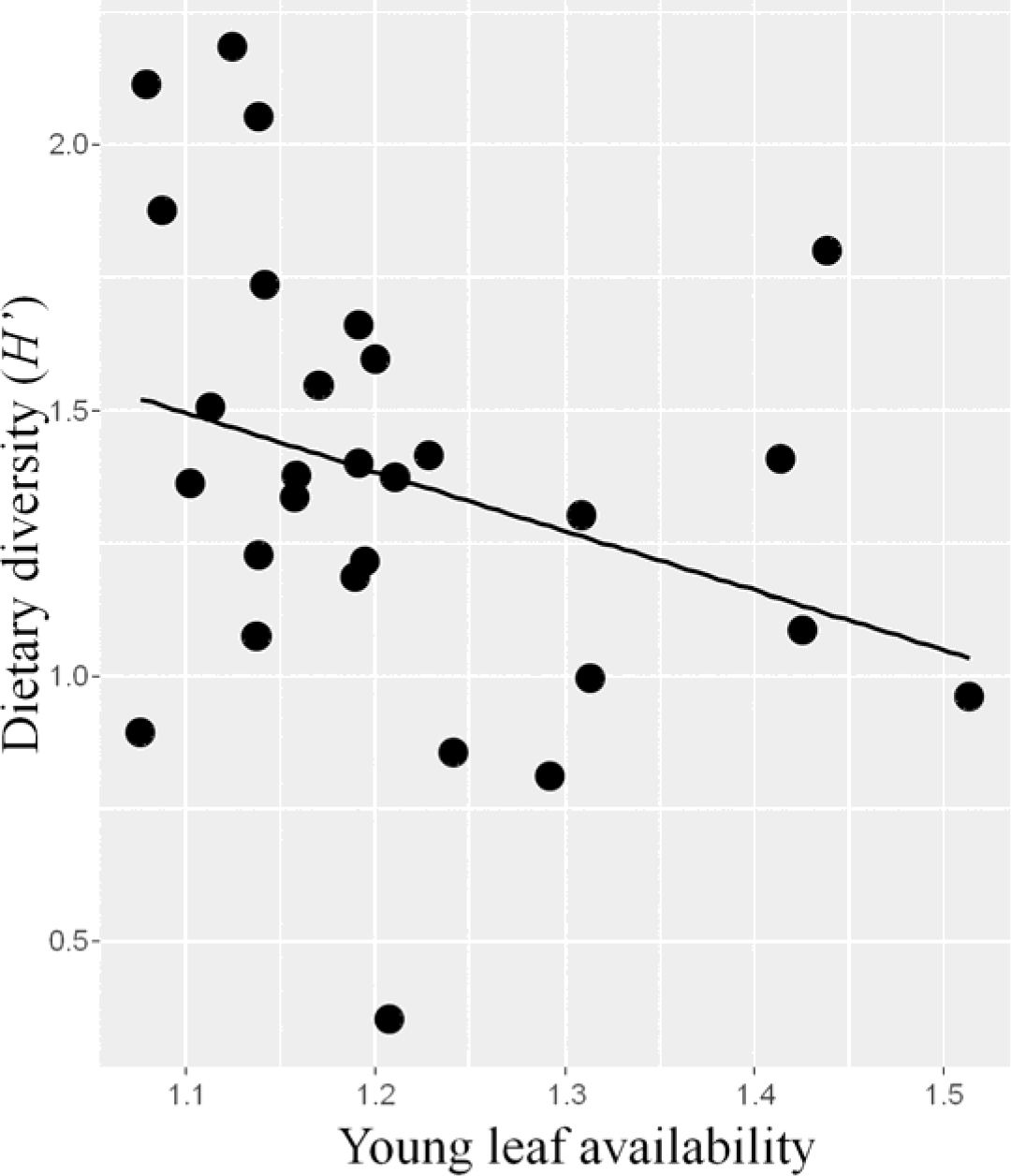
Relationship between the monthly availability of young leaves and dietary diversity.

## Discussion

In accordance with observations of previous studies on feeding behaviors of guerezas, we confirmed a high level of leaf-eating activity in the study group in general: Kalinzu Forest, Uganda, 87% (this study); Kibale National Park, Uganda, 54%–94% (Harris and Chapman 2007; Oates 1977; Wasserman and Chapman 2003); Borena–Sayint National Park, Ethiopia: 73% (Hussein et al. 2017); and Ituri, DR Congo, 58% (Bocian 1997, but refer to Fashing 2011). However, this dietary pattern is contrary to those of other guereza populations that have been observed, such as a relatively low level of leaf-eating activity but a higher level of fruit-eating activity in Kakamega, Kenya (leaves, 48%–57%; fruits/seeds, 33%–44%) (Fashing 2001) and Dja Faunal Reserve, Cameroon (leaves, 36%; fruits, 55%) (Poulsen et al. 2002). Available dietary data indicate that the guereza, which is the most widely distributed species of the genus in Africa, is ecologically flexible and adapts to different local environmental conditions. Indeed, during the study period, fruit-eating activity accounted for 45% of monthly feeding records, indicating their dietary flexibility even in response to seasonal conditions.

Moreover, we observed that the consumption of fruits and flowers in the study group significantly increased with the decrease in availability of leaf, but their fruit consumption did not significantly increase when fruit availability was high, suggesting that the tendency of choosing fruits is not strong. Additionally, the study group relied on the young leaves of *C. durandii* and fed heavily on them throughout the study period. Since the significant correlations between the leaf-eating of *C. durandii* and monthly leaf, fruit and flower availability were not detected, the operational definition of fallback foods, i.e., foods whose consumption is negatively correlated with the availability of preferred foods (Lambert and Rothman 2015; Marshall and Wrangham 2007), does not apply to this case. Nonetheless, a tendency shared by many groups of guerezas to heavily consume *C. durandii* leaves has been reported in Kibale National Park, Uganda, because of the nutrients in *C. durandii* leaves. These leaves are considered high-quality because of their high protein-to-fiber ratio (Chapman et al. 2003; Harris and Chapman 2007). Considering that they are nutrient-dense and are relatively abundant (easy to find and access), the study group may exhibit a preference for *C. durandii* leaves. Further evaluation of their dietary choices in terms of chemical and physical properties, depending on their digestive physiology and the availability of food resources (e.g., Chapman and Chapman 2002; Matsuda et al. 2017), is necessary in future studies.

From a range of 28 to 43 different food species that were reported to be consumed by other populations at Kakamega, Kibale National Park, and Ituri, the study group consumed a total of 31 food species (39 if we include the preliminary observation), which was within the specified range. However, the mean monthly *H’* of the study group (1.37) was lower than that of other populations in Kibale National Park (1.72), Ituri (1.90) and Kakamega (1.61 and 1.73), as compared with that reported by Fashing (2001). Considering the fact that the total number of plant species used as a food source by the study group was much lower than that described in the vegetation survey (68 species), we support Fashing (2001)’s statement that “guerezas appear to be adapted to feed on relatively few food species and to maintain a low dietary species diversity even in species-rich rain forest environments.”

Monthly dietary diversity increased as the number of available plants with young leaves decreased; it was noted that the effects of the availability of fruits and flowers were insignificant (Table 3). Much of the dietary diversity in the study group is seemingly attributable to the young leaf portion of their diet. Because the overall availability of young leaves declined during the study period, the guerezas were possibly forced to consume different species. This tendency of increased dietary diversity during periods when high-quality foods (i.e., young leaves and/or fruits) are scarce has also been reported in several colobine species like *C. guereza* (Fashing 2001), *Nasalis larvatus* (Matsuda et al. 2009; Yeager 1989), *Presbytis entellus* (Newton 1992), *P. rubicunda* (Davies 1991) and *Trachypithecus francoisi* (Zhou et al. 2006). Such relationships between dietary diversity and plant abundance/availability appear to be one of the general survival strategies of colobines.

In conclusion, the study group of guerezas in the Kalinzu Forest was highly folivorous and relied heavily on *C. durandii* leaves, possibly because of the high amounts these leaves being nutrient-dense and highly abundant in the study site, as observed elsewhere in Uganda. However, the study group also showed an increase (45%) in the consumption of seeds and whole fruits in response to local environmental conditions, i.e., when the leaf availability decreased, indicating the dietary flexibility of guerezas. Given that we studied a single group during one time period, this study may not be representative of the typical dietary patterns of a species or population, even at a single site. However, though our detailed descriptive study based on the behavioural observation over 30 months would be useful to provide data for further comparative meta-analysis to understand the ecological flexibility of this species in different habitats.

## Acknowledgements

We are grateful to the Uganda National Council for Science and Technology, the Uganda Forestry Department, and Uganda Wildlife Authority for permission to work in the Kalinzu Forest. Our appreciation goes to the research assistants and managers from this project, particularly, Mina Isaji, Hodaka Matsuo, Natsumi Aruga, Reiko Okano and Charles Lakwo because, without their a lot of assistances for accommodating and facilitating data collection for this project, it would not have been possible to conduct this study. We also thank Olivia Wanyana Maganyi of the Herbarium of Makerere University for their help in identification of plants and Yuri Oi and Nao Oi for their help in data entry work. This study was partly financed by the HOPE and Human Evolution Project of the Primate Research Institute, Kyoto University; JSPS KAKENHI (#19H03308 to IM, #25304019 to CH, #16H02753 to TY, #21255006 and #25257409 to HI, #24570257 to YT) and Strategic Young Researcher Overseas Visits Program for Accelerating Brain Circulation from JSPS (to H Hirai). All research was conducted in compliance with animal care regulations and applicable Uganda laws

